# MreC and MreD balance the interaction between the elongasome proteins PBP2 and RodA

**DOI:** 10.1101/769984

**Authors:** Xiaolong Liu, Jacob Biboy, Waldemar Vollmer, Tanneke den Blaauwen

## Abstract

Rod-shape of most bacteria is maintained by the elongasome, which mediates the synthesis and insertion of peptidoglycan into the cylindrical part of the cell wall. The elongasome contains several essential proteins, such as RodA, PBP2, and the MreBCD proteins, but how its activities are regulated remains poorly understood. Using *E. coli* as a model system, we investigated the interactions between core elongasome proteins *in vivo*. Our results show that PBP2 and RodA form a complex mediated by their transmembrane and periplasmic parts and independent of their catalytic activity. MreC and MreD also interact directly with PBP2. MreC elicits a chance in the interaction between PBP2 and RodA, which is suppressed by MreD. The cytoplasmic domain of PBP2 is required for this suppression. We hypothesize that the *in vivo* measured PBP2-RodA interaction change induced by MreC corresponds to the conformational change in PBP2 as observed in the MreC-PBP2 crystal structure, which was suggested to be the “on state” of PBP2. Our results indicate that the balance between MreC and MreD determines the activity of PBP2, which could open new strategies for antibiotic drug development.

**Importance:** The cell envelope of *Escherichia coli* bears the protective and shape-determining peptidoglycan layer sandwiched between the outer and inner membranes. Length growth in bacteria is accomplished by a protein complex termed elongasome. We used Förster Resonance Energy Transfer (FRET) that reports not only on whether proteins interact with each other but also on conformational changes during interactions, to investigate how the elongasome might be activated. RodA and PBP2 provide the peptidoglycan glycosyltransferase and transpeptidase activities needed to synthesize new peptidolgycan during length growth, respectively, and PBP2 activates RodA. We show that the interactions between MreC and MreD with PBP2-RodA alter the nature of the interaction between PBP2 and RodA and hypothesis that the corresponding conformational change in the PBP2-RodA complex allows switching between the ‘on’ and ‘off’ states of the elongasome.

## Introduction

Bacterial cells are surrounded by a peptidoglycan layer that maintains their shape and protects them from bursting due to the osmotic pressure. The biosynthesis of peptidoglycan is the target of many antibiotics that are used in clinical therapies for bacterial infections. The spread of antibiotic resistant pathogens calls urgently for the development of novel antibiotics. In depth knowledge on peptidoglycan synthesis will aid in the development of effective screening assays to select cell wall synthesis inhibitors. Peptidoglycan is a mesh-like heteropolymer of glycan chains of Glc*N*Ac-Mur*N*Ac-peptide subunits that are connected by peptide cross-links (1). Peptidoglycan synthesis begins in the cytoplasm with synthesis of UDP-Glc*N*Ac and UDP-Mur*N*Ac-pentapeptide (2). Two following membrane steps, catalyzed by MraY and MurG, assemble the precursor lipid II (3, 4), which is flipped to the periplasmic side of the cytoplasmic membrane by lipid II flippase(s) MurJ and/or FtsW (5–7). Glc*N*Ac-Mur*N*Ac pentapeptide units are polymerized into glycan chains and the peptides are cross-linked to bridge the glycan stands by peptidoglycan synthases to expand the peptidoglycan layer while the carrier lipid is recycled (8–10). Most rod-shaped bacteria employ two protein complexes, elongasome and divisome, to guide peptidoglycan synthesis during lateral growth and cell division, respectively (11).

In *E. coli*, the divisome contains more than twenty proteins. Assembly of the divisome starts with positioning the FtsZ ring at midcell together with other early divisome proteins, such as FtsA, ZipA, ZapA and FtsEX, to form the early divisome (12–14) Subsequently, the late divisome proteins, FtsK, FtsBLQ, FtsW, PBP3 and FtsN, are recruited (15). These proteins localize to midcell in an interdependent order (15, 16). Among these proteins, FstW, PBP3 and PBP1B provide the peptidoglycan synthesis activity during septum synthesis (10, 17, 18). PBP1B has both glycosyltransferase (GTase) and transpeptidase (TPase) activity (19), while FtsW and PBP3 (also called FtsI) only have GTase activity and TPase activity, respectively (10). Although the mechanisms of peptidoglycan synthesis regulation is not fully understood, recent studies showed that the cell division proteins have competing effects and either inhibit (FtsQLB complex) or stimulate the activities of FtsW-PBP3-PBP1B (20–22).

Proteins that are known to be part of the elongasome are the cytoplasmic membrane associated actin homologue MreB, the bitopic membrane proteins RodZ, MreC and PBP2, and the integral membrane proteins MreD and RodA (Fig. 1a). MreB polymerizes into short filaments that rotate around the cylindrical part of the cell (23, 24). The rotation of MreB is believed to drive the topography of the insertion of peptidoglycan into the lateral wall (23, 25–27). Bacterial two hybrid analysis showed that MreB interacts with MreC, but not with MreD (28), while RodZ interacts strongly with itself, MreB and MreC (28, 29) (Fig.1b), and these interactions are essential to maintain bacterial morphology (28, 30–32). RodA and PBP2 form a stable subcomplex (33) and provide GTase and TPase activity, respectively, during cylindrical peptidoglycan synthesis (9, 34, 35). This subcomplex also shows a circumferential motion that is similar to that of MreB. The bifunctional GTase-TPase PBP1A interacts with PBP2 and stimulates its activity (18). Because PBP1A moves independently of the rotation of PBP2 and MreB, it is thought not to be part of the core elongasome (18, 36). However, the function and role of most elongasome proteins are still poorly understood. How peptidoglycan synthesis is activated and regulated during elongation is still the key question. In this study, combining genetics, microscopy and Förster Resonance Energy Transfer (FRET), we investigated the functions of, and interactions between, these core elongasome proteins. The transfer of energy between a donor fluorescent-protein fusion and an acceptor fluorescent-protein fusion (FRET) is very sensitive to distance, which even allows the detection of conformational changes that affect this distance (7). Our results indicate that MreC and MreD modulate the interaction between PBP2 and RodA in oppositely, which likely reflects a mechanism of elongasome activation and regulation.

**Figure 1.**
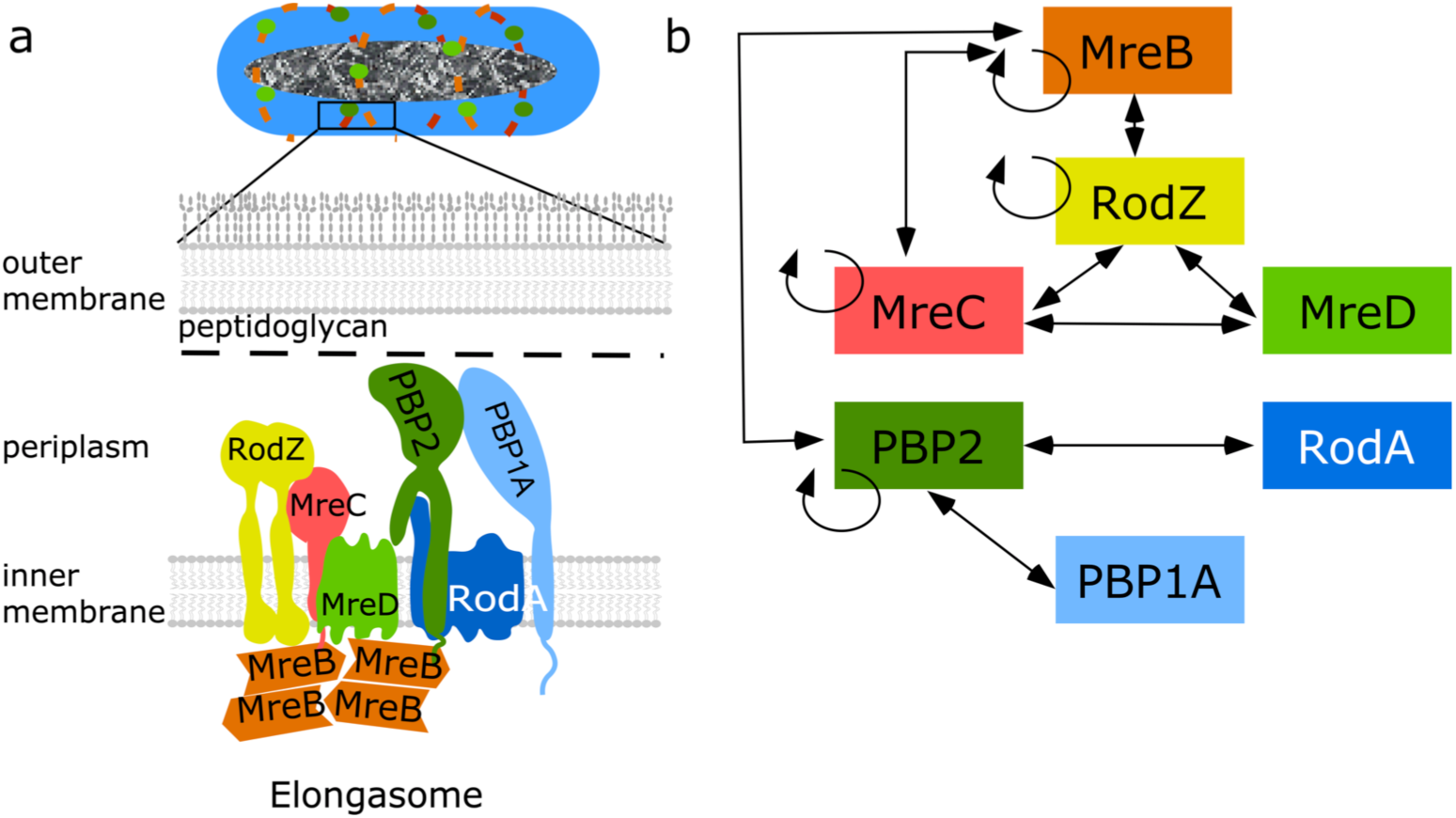
Core elongasome proteins and their interactions in *E. coli*. **a.** Schematic representation of the *E. coli* cell envelope and elongasome. MreB localizes in patches underneath the cytoplasmic membrane and recruits other elongasome proteins. The peptidoglycan layer is sandwiched by the cytoplasmic membrane and the outer membrane. **b**. Identified interactions between elongasome proteins from previous interaction studies (1–5). Double arrowed lines represent the interaction between different proteins. Circular arrows indicate self-interaction.

## Results

### RodA and PBP2 activities are not essential for their interaction

RodA and PBP2 form a stable peptidoglycan synthesizing subcomplex in the cytoplasmic membrane as detected by FRET (33). To investigate whether this interaction relies on their enzymatic activities, RodA^R109A^ and RodA^Q207R^ versions, which were predicted to be inactive based on studies on its homologue FtsW, were constructed (Supplementary Fig. 1) (5, 38). As expected, these mutants could not complement the temperature sensitive RodA strain LMC882 at the non-permissive temperature, and the RodA^Q207R^ variant even showed dominant negative effects at the permissive temperature (Fig. 2a). Subsequently, N-terminal mCherry fused versions (33, 34) of the inactive RodA proteins were expressed to test their interaction with mKO-PBP2^WT^ by FRET (Fig. 2b). In our FRET system, the direct fused mCherry-mKO tandem was used as positive control (33). To account for possible interactions between proteins due to crowding in the cytoplasmic membrane, an integral membrane protein unrelated to peptidoglycan synthesis, GlpT^3,34^, was fused to mKO, and its interaction with mCh-RodA was detected as negative control. The acceptor FRET efficiency values (Ef*_A_*) of all FRET samples were calculated using our previously published mKO-mCh FRET spectral unmixing method (33) (Supplementary Fig. 2a). An Ef*_A_* value of 31.0 ± 4.0% was observed for the tandem control (Fig. 2b and Table 1), which is comparable to the published data (33). An Ef*_A_* value of 1.1 ± 3.5% was observed for the RodA-GlpT negative control (Fig. 2b and Table 1). FRET experiments with PBP2^WT^ and RodA^R109A^ or RodA^Q207R^ yielded Ef*_A_* values of 12.5 ± 1.9% and 12.7 ± 1.2%, respectively, which are comparable to the Ef*_A_* value of 12.7 ± 1.7% of wild type RodA, indicating an interaction between PBP2^WT^ and all RodA versions (Fig. 2b, Supplementary Fig 2a and Table 1). To determine whether the activity of PBP2 was required for the interaction with RodA, we expressed the inactive variant PBP2^S330C^, which is not able to bind benzylpencillin (35). PBP2^S330C^ had a strong dominant negative effect during the complementation in the PBP2 temperature sensitive strain LMC582 (Fig. 2c), while the detected Ef*_A_* value of PBP2^S330C^-RodA^WT^ remained 10.9 ± 0.5%, which was slightly below the Ef*_A_* value of PBP2^WT^-RodA^WT^ (Fig. 2b and Table 1). These results imply that the activities of RodA and PBP2 are not needed for their interaction.

**Figure 2.**
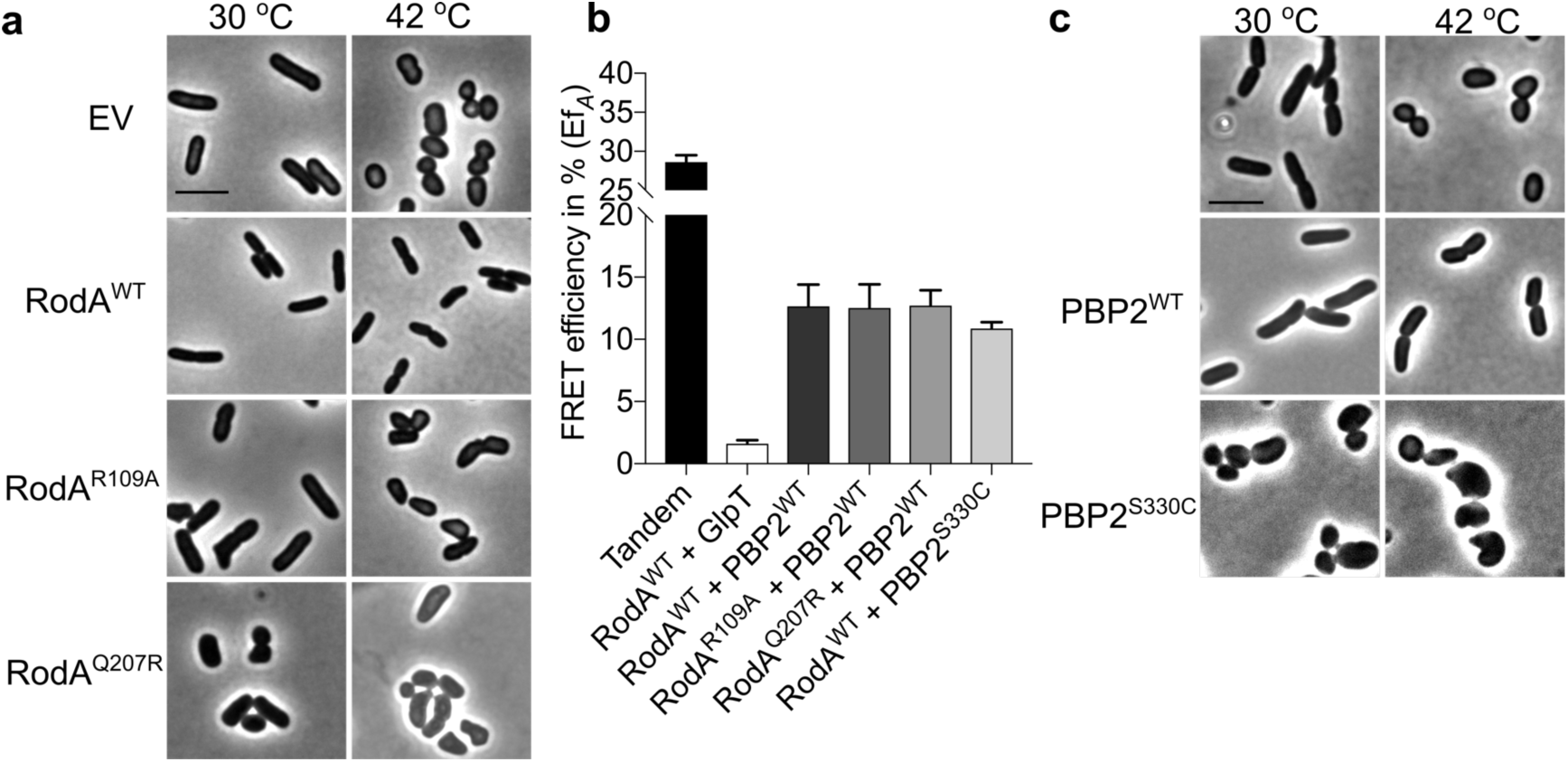
Activity of RodA and PBP2 are not required for their interaction. **a,** Phase contrast images of the complementation of RodA variants. RodA temperature sensitive strain LMC882 was transformed with plasmids expressing RodA variants and grown in LB medium at 30 °C (left panel) and 42 °C (right panel) for 2 mass doublings (with15 μM IPTG induction). EV, empty vector. **b**, Calculated acceptor FRET efficiencies (Ef*_A_*) between PBP2 and RodA variants from spectral FRET measurements. RodA and its variants are fused with mCherry. PBP2 and its variants are fused with mKO. **c**. Phase contrast images of the complementation of PBP2 variants. The PBP2 temperature sensitive strain LMC582 was transformed with PBP2^WT^ or PBP2^S330C^, and grown in LB medium at 30 °C (left panel) and 42 °C (right panel) for 2 mass doublings (with 15 μM IPTG induction). Scale bar equals 5 μm. All the results in the figure are representative of at least three independent experiments.

**Table 1.** Summary of the calculated acceptor FRET efficiencies (Ef*_A_*) from spectral FRET measurements for listed samples.

| Parameter | Proteins expressed |  | Ef<sub>A</sub> (%) | SD (%) | N<sup>1</sup> |
| --- | --- | --- | --- | --- | --- |
|  | pTHV037 | pSAV057 |  |  |  |
| **Positive control<sup>2</sup>** |  |  |  |  |  |
| Tandem | Empty plasmid | mKO-mCh | 31.0 | 4.0 | 22 |
| **Negative control<sup>3</sup>** |  |  |  |  |  |
| RodA-GlpT | mCh-RodA | mKO-(GGS)<sub>2</sub>-GlpT | 1.1 | 3.5 | 24 |
| **Biological interactions** |  |  |  |  |  |
| RodA<sup>4</sup>-PBP2<sup>WT</sup> | mCh-RodA | mKO-PBP2<sup>WT</sup> | 12.7 | 1.7 | 16 |
| RodA<sup>R109A</sup>-PBP2 | mCh-RodA<sup>R109A</sup> | mKO-PBP2<sup>WT</sup> | 12.5 | 1.9 | 4 |
| RodA<sup>Q207R</sup>-PBP2 | mCh-RodA<sup>Q207R</sup> | mKO-PBP2<sup>WT</sup> | 12.7 | 1.2 | 4 |
| RodA-PBP2<sup>S330C</sup> | mCh-RodA | mKO-PBP2<sup>S330C</sup> | 10.9 | 0.5 | 3 |
| RodA-MalFNT-PBP2 | mCh-RodA | mKO-MalFNT-PBP2 | 14.4 | 1.7 | 4 |
| RodA-MalF37-PBP2 | mCh-RodA | mKO-MalF37-PBP2 | 8.2 | 1.3 | 3 |
| RodA-PBP2<sup>L61R</sup> | mCh-RodA | mKO-PBP2<sup>L61R</sup> | 12.8 | 2.8 | 8 |
| RodA-RodA<sup>WT</sup> | mCh-RodA | mKO-RodA | 5.5 | 1.4 | 4 |
| MreB<sup>SW</sup>-RodA | mCh-MreB<sup>SW5</sup> | mKO-RodA | 7.2 | 3.4 | 4 |
| PBP2<sup>WT</sup>-PBP2<sup>WT</sup> | mCh-PBP2<sup>WT</sup> | mKO-PBP2 | 9.2 | 0.5 | 4 |
| PBP2-MalFNT-PBP2 | mCh-PBP2<sup>WT</sup> | mKO-MalFNT-PBP2 | 10.5 | 2.0 | 4 |
| MreC-PBP2<sup>WT</sup> | mCh-MreC | mKO-PBP2<sup>WT</sup> | 5.1 | 1.2 | 6 |
| MreC-MalFNT-PBP2 | mCh-MreC | mKO-MalFNT-PBP2 | 5.4 | 1.7 | 6 |
| MreC-PBP2<sup>L61R</sup> | mCh-MreC | mKO-PBP2<sup>L61R</sup> | 5.3 | 0.6 | 2 |
| MreD-PBP2<sup>WT</sup> | mCh-MreD | mKO-PBP2<sup>WT</sup> | 4.3 | 1.2 | 10 |
| MreD-MalFNT-PBP2 | mCh-MreD | mKO-MalFNT-PBP2 | 5.3 | 1.6 | 10 |
| MreCD<sup>6</sup>-PBP2<sup>WT</sup> | mCh-MreCD | mKO-PBP2<sup>WT</sup> | 3.3 | 0.5 | 6 |
| MreCD<sup>7</sup>-MalFNT-PBP2 | mCh-MreCD | mKO-MalFNT-PBP2 | 5.4 | 1.1 | 5 |
| **+ mecillinam** |  |  |  |  |  |
| **Positive control** |  |  |  |  |  |
| Tandem | Empty vector | mKO-mCh | 31.0 | 0.6 | 2 |
| **Negative control** |  |  |  |  |  |
| RodA-GlpT | mCh-RodA | mKO-(GGS)<sub>2</sub>-GlpT | 3.1 | 0.1 | 2 |
| **Biological interactions** |  |  |  |  |  |
| RodA-PBP2<sup>WT</sup> | mCh-RodA | mKO-PBP2<sup>WT</sup> | 8.6 | 1.1 | 6 |
| RodA-MalF37-PBP2 | mCh-RodA | mKO-MalF37-PBP2 | 3.0 | 2.0 | 4 |
| RodA-PBP2<sup>S330C</sup> | mCh-RodA | mKO-PBP2<sup>S330C</sup> | 7.9 | 1.0 | 2 |
| RodA-PBP2<sup>L61R</sup> | mCh-RodA | mKO-PBP2<sup>L61R</sup> | 9.7 | 2.2 | 3 |

<sup>1</sup> No, number of biological repeats;
<sup>2,3</sup> Here all measured positive and negative controls are averaged. In the figures the controls are included that belong to the corresponding measurements.
<sup>4</sup> RodA without superscript represent the wild type version.
<sup>5</sup> mCh-MreB<sup>SW</sup> is a sandwich fusion of MreB-mCherry-MreB (33).
<sup>6,7</sup> MreC and MreD were expressed from one plasmid, and MreC was fused to mCherry while MreD was non-fused.

### The transmembrane and periplasmic parts of PBP2 contribute to its interaction with RodA

To reveal which part of PBP2 interacts with RodA, two domain swap mutants of PBP2 were constructed. The cytoplasmic N-terminus (NT) and or the N-terminal region with the transmembrane-helix (NT-TMH) of PBP2 were replaced by the corresponding N-terminal stretches of MalF, a bitopic membrane protein that has been previously used in domain swap studies (39–41), to yield ^MalFNT^PBP2 and ^MalF37^PBP2, respectively (Fig. 3a). Both versions of PBP2 were able to localize to the membrane but showed dominant negative effects, indicating the essentiality of the replaced parts (Fig. 3b). The replacement of the NT of PBP2 did not change its interaction with RodA, as the detected Ef*_A_* value remained 14.4 ± 1.1%, which was not significantly different compared to that of the interaction between RodA and wild type PBP2 (Fig. 3c, Table 1 and Supplementary Fig. 2b). However, replacement of the TMH of PBP2 significantly reduced the Ef*_A_* value between PBP2 and RodA to 8.2 ± 1.3%, which reflected an apparent distance increase from 8.6 nm to 9.8 nm between the two proteins (42) (Fig. 3c and Table 1). This decrease in distance was not caused by a change in amount of measured fluorescence for the RodA and PBP2 fusions expressed in the cells (Supplementary Fig. 2b), or due to the shorter transmembrane helix after replacement (Fig. 3a). The average rise per residue in transmembrane helices is 0.15 nm (43), therefore the two amino acid residues shorter helix in ^MalF37^PBP2 (Fig. 3a) could maximally change the distance between donor and acceptor fluorophores by 0.3 nm. The still considerably higher Ef*_A_* value compared to the negative control indicates that the transmembrane helix alone is not sufficient for the interaction between PBP2 and RodA and that the periplasmic domain of PBP2 is also involved in this interaction (Fig. 3d).

**Figure 3.**
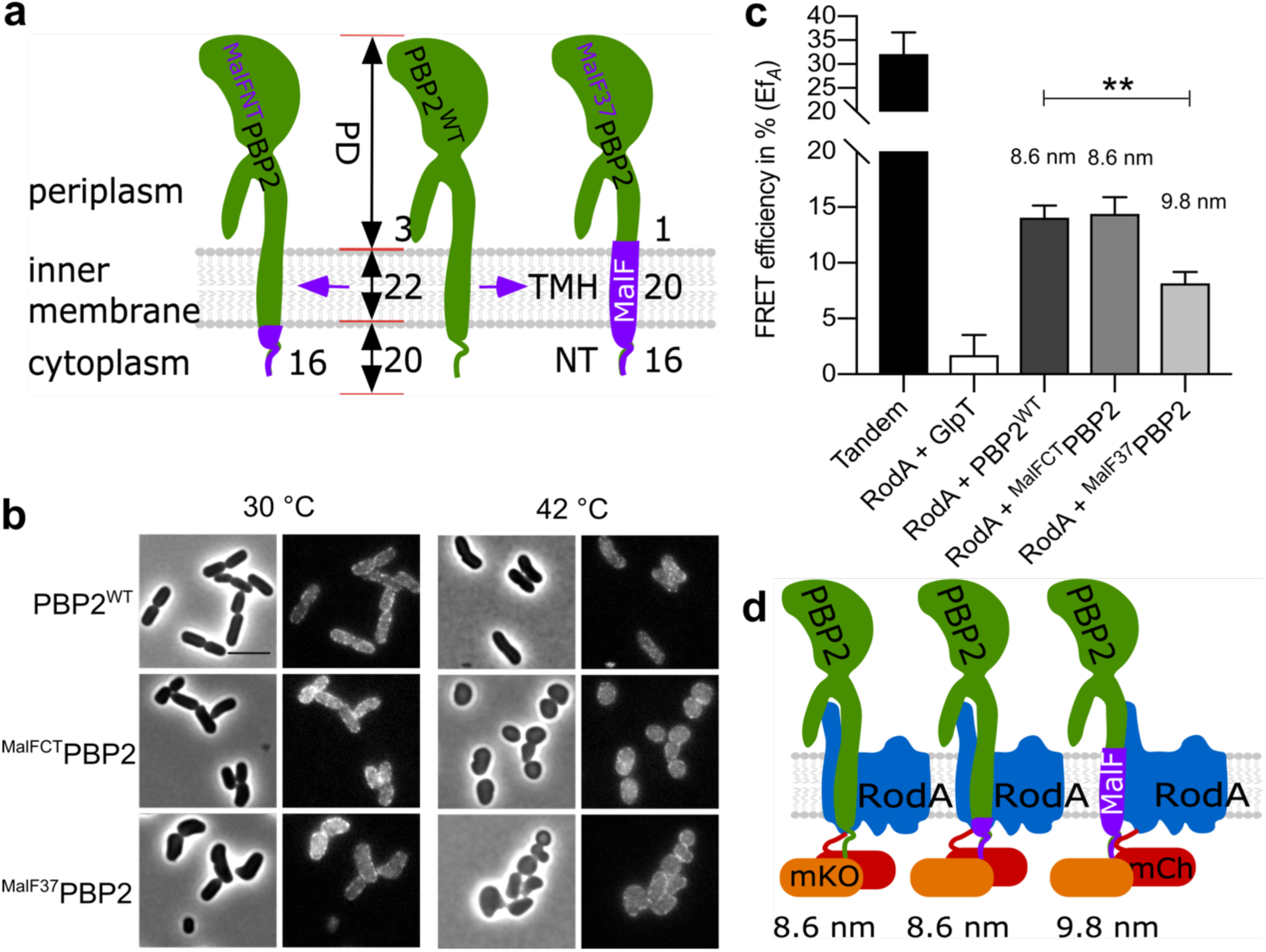
Functionality and interaction of PBP2 domain swap mutants. **a.** Schematic illustration of PBP2 domain-swap mutants. NT: N-terminus; TMH: transmembrane helix; PD: periplasmic domain; PBP2^WT^: wild type PBP2; ^MalFNT^PBP2: the cytoplasmic N-terminus of PBP2 was replaced with the MalF cytoplasmic N-terminus; ^MalF37^PBP2: the NT and TMH domains of PBP2 were replaced with corresponding domains of MalF. Numbers indicate the residues involved in replacements in each domain of the proteins. **b**. Phase contrast and fluorescence images of the complementation of PBP2 mutants. The PBP2 temperature sensitive strain LMC582 was transformed with the PBP2 variants, and grown in LB medium at 30 °C (left panels) and 42 °C (right panels) for 2 mass doublings (with 15 μM IPTG induction). Scale bar equals 5 μm. **c**. Calculated acceptor FRET efficiencies (Ef*_A_*) between PBP2 and RodA variants from spectral FRET measurements. RodA and its variants are fused with mCherry. PBP2 and its variants are fused with mKO. P value determined with Student’s t-test (**: p<0.001). The numbers are apparent distances between the two proteins (fluorophores) and were calculated using the equation, E = (1+(r/R_0_)^6^)^-1^, where r is the distance between the chromophores and R_0_ (the Förster distance) is 6.4 nm for the mCh-mKO pair. **d**. Schematic illustration of the interaction between RodA and PBP2 variants. After replacement of the TMH domain of PBP2, by the TMH of MalF, the FRET efficiency is lower indicating that the distance between the FP fused TMHs of PBP2 and RodA has increased. However the interaction is not lost, indicating that periplasmic domains are also involved in the interaction All results in the figure are representative of at least three independent experiments.

### MreC interacts with PBP2 and affects PBP2-RodA interaction

A recent study of PBP2-MreC from *Helicobacter pylori* showed two different structural conformations of PBP2 in the MreC-bound and unbound forms, respectively^43^ (Fig. 4 a). The authors proposed that the binding of MreC to the periplasmic hydrophobic zipper of PBP2 induces a conformational change in PBP2 and a switch from an off state into an on state (44). In our FRET system about 1000 copies of the mKO fusion proteins per cell are expressed from a plasmid (45). The ∼180 endogenous copies of MreC molecules (46) are not sufficient to activate the majority of the by plasmid expressed mKO-PBP2 molecules. Therefore, we hypothesize that most of the mildly overexpressed PBP2 versions remain in the off state conformation (Fig. 4b, left). We reasoned that the interaction between PBP2 and RodA could be sensitive to possible conformational changes of PBP2, if we would additionally express MreC to balance the molecule numbers of both proteins. To this end we first tested the interaction between a functional mCh-MreC (28) fusion and mKO-PBP2 by FRET measurements. The observed Ef*_A_* value of 5.1 ± 1.2% indicates a direct interaction between PBP2 and MreC (Fig. 4c, table 1 and Supplementary Fig. 3), which is in agreement with the structural study of PBP2-MreC from *Helicobacter pylori* (44).

**Figure 4.**
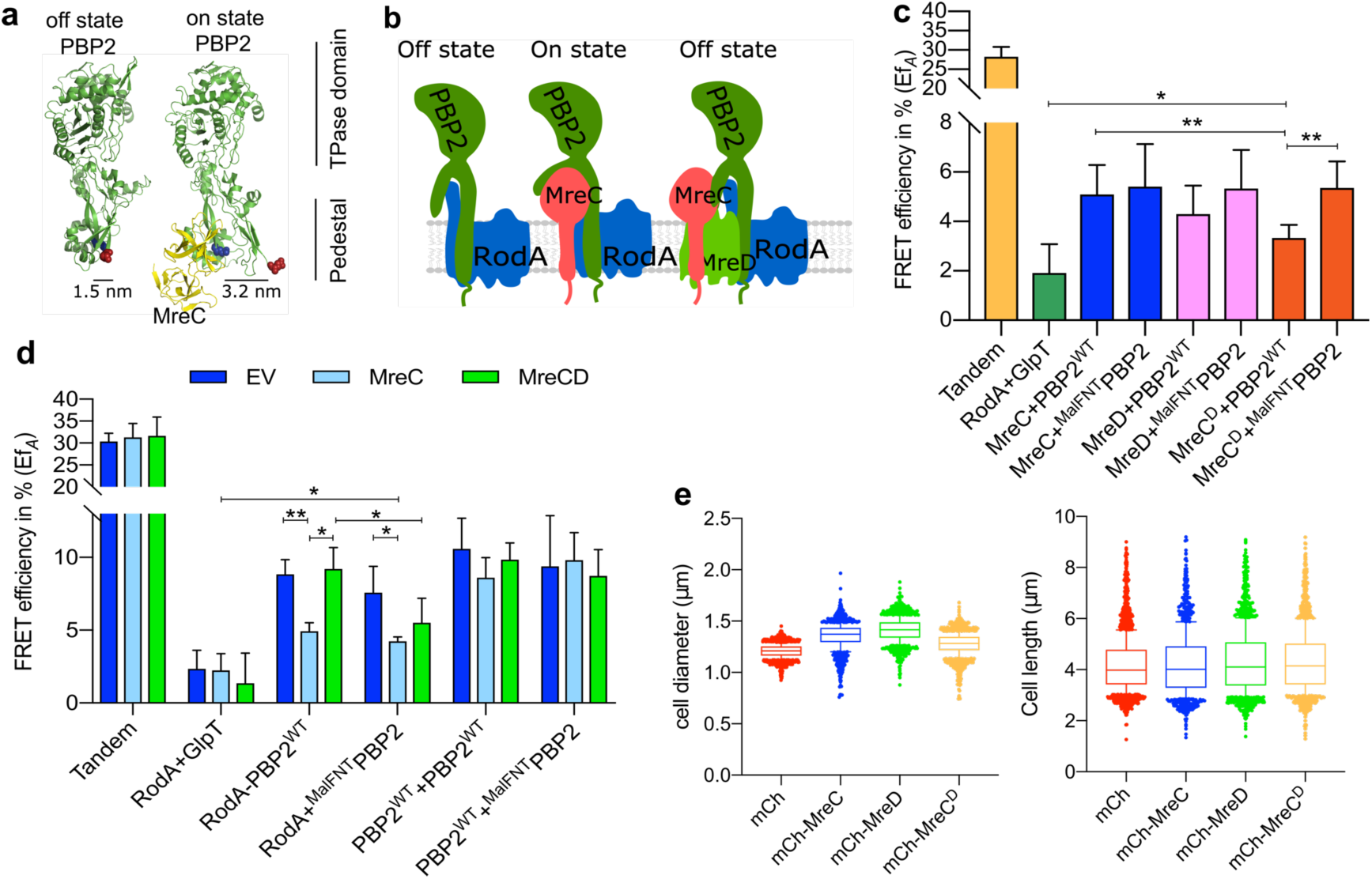
The balance of MreC and MreD affects the interaction between RodA and PBP2. **a.** Crystal structures of *E. coli* PBP2 in different conformations (6) were modeled from *Helicobacter pylori* PBP2 structures (7) using Phyre2 (8). The structural information lacks the juxta-membrane, transmembrane helix and cytoplasmic regions of PBP2. MreC binds to PBP2 and was proposed to switch PBP2 from the “off state” to the “on state” (7). The distances between the two sphered residues, (blue for Glu157) and (red for Lys60) were calculated based on the structure of PBP2 in the different conformations. **b**. Schematic representation of PBP2 conformational changes caused by MreC. Left panels: PBP2 stays in the “off state” in the absence of MreC (the distance between the cytoplasmic terminus of RodA and PBP2 is small); middle panels: PBP2 switches to the “on state” after binding MreC (the distance between cytoplasmic terminus of RodA and PBP2 is larger), right panels: MreD suppresses the MreC-mediated conformational change of PBP2 and keeps PBP2 in the “off state”. **c**. Calculated acceptor FRET efficiencies (Ef*_A_*) between MreCD proteins and RodA variants from spectral FRET measurements. MreC: mCherry fused MreC; MreD: mCherry fused MreD; MreC^D^: MreCD co-expressed from the same plasmid, and MreC is fused with mCherry while MreD is non-fused. PBP2 and its variants are fused with mKO. **d**. Calculated acceptor FRET efficiencies (Ef*_A_*) between RodA and PBP2 variants from spectral FRET measurements in the three-plasmids FRET experiments. EV: a third empty vector; MreC: a third plasmid expressing non-fused MreC; MreCD: a third plasmid expressing non-fused MreCD. **e**. Cell length and diameter changes after expressing MreC, MreD or MreCD together. LMC500 strain was transformed with each construct and grown in LB medium at 37 °C and induced with 15 μM IPTG for 2 mass doublings. Proteins were expressed from the pSAV057 derived plasmids (mKO: control; MreC: mKO fused MreC; MreD: mKO fused MreD; MreC^D^: co-expression of MreC and MreD, MreC is fused with mKO while MreD is not fused. About 1000 cells were analyzed. P value determined with Student’s t-test (*: p<0.05,; **: p<0.01:).

We next we employed a three-plasmids-FRET system that expressed MreC from a third plasmid when testing the interaction between PBP2 and RodA (Fig. 4d, Table 2 and Supplementary Fig. 4). A control strain contained an empty plasmid instead of the MreC-expression plasmid. In the presence of the empty plasmid, the calculated Ef*_A_* values for the tandem (positive control) and RodA-GlpT (negative control) were 30.4 ± 1.8% and 2.3 ± 1.3%, respectively (Fig. 4d and Table 2). These Ef*_A_* values remained unchanged in the presence of MreC expressed from the third plasmid (Fig. 4d and Table 2). Interestingly, the Ef*_A_* value for the RodA-PBP2 interaction was significantly reduced to 4.9 ± 0.6% in the presence of MreC, compared with the Ef*_A_* of 8.8 ± 1.1% in the presence of empty plasmid (Fig. 4d and Table 2). These results indicate that MreC changes the interaction between PBP2 and RodA, which would be consistent with an conformational change of PBP2 from the off state to the on state proposed from the crystal structures (44) (Fig. 4b, middle).

**Table 2.**
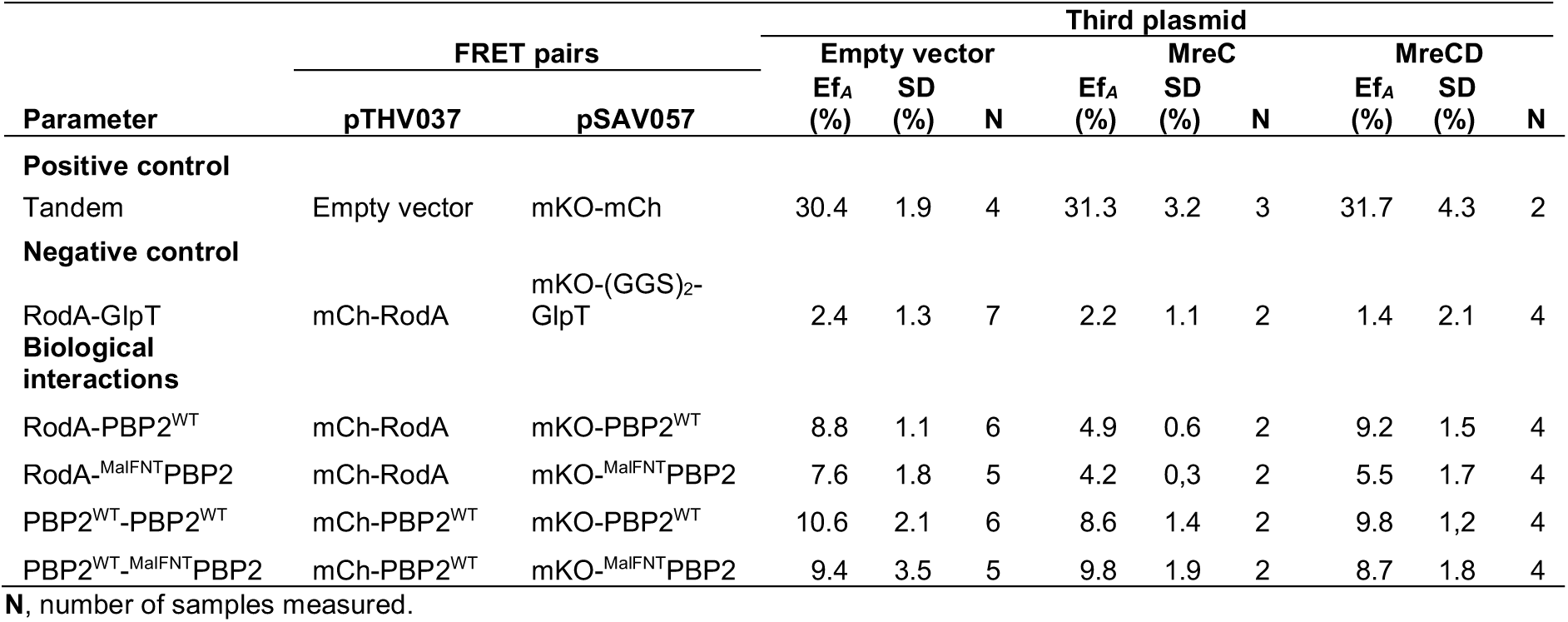
Summary of the calculated acceptor FRET (Ef*_A_*) efficiencies from spectral FRET measurements for listed samples.

### MreD suppresses the MreC-mediated change in the PBP2-RodA interaction

During our study, we noticed that overexpression of MreC caused morphological defects of the wild type strain, increasing the diameter of *E. coli* cells (Fig. 4e and Supplementary Fig. 5). Interestingly, the co-expression of MreD together with MreC suppressed these morphological defects and restored the wild type phenotype (Fig. 4e and Supplementary Fig. 5). To further investigate this effect, we aimed to express an N-terminal functional mCherry fusion of MreD (30) which is an integral membrane protein with 6 predicted transmembrane helices and both termini localized in the cytoplasm (Supplementary Fig. 6). Consistent with this topology model we readily observed fluorescence signals for N-terminal fused GFP-MreD (30) and mKO-MreD versions, which could not be possible if the N-terminus of MreD would localize in the oxidative periplasm where GFP and mKO do not mature (37). Similarly as for MreC, the overexpression of MreD alone resulted in morphological defects of *E. coli* cells (Fig. 4e and supplementary Fig. 5).

To study the role of MreD in the elongasome, FRET experiments were designed to detect a possible interaction with PBP2. The Ef*_A_* value of 4.3 ± 1.1% indicated a direct interaction between MreD and PBP2 (Fig. 4c, Table 1 and Supplementary Fig. 3). Subsequently, the interaction between MreC and PBP2 was measured by FRET in the presence of MreD. The calculated Ef*_A_* between MreC and PBP2 was significantly reduced from 5.1 ± 1.2% to 3.3 ± 0.5% (p=0.0078) when MreD was co-expressed (Fig. 4c, Table 1 and Supplementary Fig. 3). Since MreC reduced the Ef*_A_* of RodA-PBP2 from 8.8 ± 1.1% to 4.9 ± 0.5%, it was possible that MreD altered the effect of MreC on the RodA-PBP2 interaction. Therefore, the three-plasmid FRET experiment was applied to detect the interaction between RodA and PBP2 in the presence of MreCD. Interestingly, the Ef*_A_* value of RodA-PBP2 was restored to 9.2 ± 1.5%, which was comparable with the Ef*_A_* in the presence of the third, empty plasmid (Fig. 4d and Table 2). These combined results suggest a regulatory mechanism by which MreC interacts with PBP2 and changes its conformation, while MreD interacts with MreC and PBP2 to prevent this conformational change PBP2. These conformational changes could correspond the proposed on and off states of PBP2 as published (44) (Fig. 4 a and b).

### The cytoplasmic part of PBP2 is important for the interplay with the MreCD proteins

As showed before, the cytoplasmic NT part of PBP2 has an essential unknown function rather than being involved in the RodA-PBP2 interaction (Fig. 3). We considered that the NT of PBP2 might be important for its self-interaction and or interactions with other partner proteins. However, the Ef*_A_* values of ^MalFNT^PBP2 with wild type PBP2, MreC and MreD were not different from those of wild type PBP2 (Fig. 4d, Supplementary Fig. 7 and Table 1). Interestingly, the Ef*_A_* value of the interaction between MreC and wild type PBP2, but not the ^MalFNT^PBP2, was reduced by the co-expression of MreD (Fig. 4d, Table 1 and Supplementary Fig. 4c). Similarly, in the three-plasmid FRET experiments, MreD was not able to suppress the MreC-mediated change in the ^MalFNT^PBP2-RodA interaction; the Ef*_A_* value of RodA-^MalFNT^PBP2 FRET remained at 5.5 ± 1.7% in presence of MreCD, rather than being restored to 9.2 ± 1.5% as in the RodA-PBP2^WT^ experiments (Fig. 4d, Table 2 and Supplementary Fig. 4c). Together, these results indicate that the cytoplasmic part of PBP2 plays an important role in the MreCD-mediated regulation of the interaction between PBP2 and RodA.

### MreCD proteins do not alter PBP2 self-interaction

So far our results has shown that MreC and MreD have opposite effects on the interaction between RodA and PBP2. In contrast, the interaction between two PBP2 molecules (17) was not significantly affected upon overexpression of MreC or MreCD, as the calculated Ef*_A_* values for the PBP2-PBP2 interaction remained unchanged compared to the values for the expression of the third empty plasmid (Fig. 4d and Table 2). RodA was also found to interact with itself (Table 1 and Supplementary Fig. 7). Likely PBP2 and RodA function as a complex of dimers, which might also allow simultaneously synthesis of multiple glycan strand as has been proposed (47–49).

### Effect of mecillinam on the interaction between RodA and PBP2

As showed above, both the transmembrane helix and periplasmic part of PBP2 contribute to its interaction with RodA (Fig. 3). The binding of MreC to the periplasmic hydrophobic zipper domain of PBP2, which presumably changes the conformation of PBP2 from off state to on state, reduces the detected Ef*_A_* between RodA and PBP2 (Fig. 4d and e). Interestingly, the PBP2 specific inhibitor mecillinam, also caused a reduction in the FRET efficiency of the interaction between RodA and PBP2. The RodA-PBP2^WT^ interaction pair yielded a reduced Ef*_A_* of 8.6 ± 1.1% in the presence of mecillinam (33), comparing with the Ef*_A_* of 12.7 ± 1.7% without mecillinam (Fig. 2, Table 1 and Supplementary Fig. 8). Since mecillinam binds specifically to the periplasmic TPase active site of PBP2, a possible explanation could be that binding of mecillinam reduces the affinity between the periplasmic parts of PBP2 and RodA, but does not interfere with the interaction between their transmembrane regions. Indeed, after replacing the transmembrane helix of PBP2 to abolish this part of the interaction (RodA-^MalF37^PBP2), mecilinam further reduced the Ef*_A_* value to 3.0 ± 2.0% (Supplementary Fig. 8 and Table 1), consistent with an almost complete loss of the interaction between RodA and ^MalF37^PBP2. That mecillinam causes disruption of only one out of two interacting regions between PBP2 and RodA is also in agreement with the observations that it does not disrupt the structure of the elongasome (50). The inactive mutant PBP2^S330C^, which cannot bind benzylpenicillin (35), still responded to mecillinam and showed a similar Ef*_A_* reduction as PBP2^WT^ (Supplementary Fig. 8 and Table 1). This suggests that although the inactive mutant PBP2^S330C^ does not bind mecillinam covalently, it still interacts with it.

### PBP2^L61R^ stays in the off state and activates RodA

A recent study reported a version of PBP2 in which Leu61 was replaced by Arg (PBP2^L61R^) that could suppress an MreC defect, and was proposed to stay in the on state conformation mimicking activation by MreC (51). If this would be the case, the RodA-PBP2^L61R^ pair would be expected to have a reduced FRET efficiency, since the MreC activated RodA-PBP2^WT^ pair resulted in a reduction in FRET efficiency (Fig 4d). Therefore, we constructed an N-terminal mKO fusion of PBP2^L61R^ to test the interactions with its partner proteins. Surprisingly, the Ef*_A_* for RodA-PBP2^L61R^ remained 12.8 ± 2.8%, which was comparable with the Ef*_A_* of RodA-PBP2^WT^ that was presumably in the off state (Fig. 5a and b, Table 1 and Supplementary Fig. 9a). Mecillinam reduces the Ef*_A_* value of the RodA-PBP2^L61R^ interaction to 9.7± 2.2%, which was also comparable with its effect on the RodA-PBP2^WT^ interaction (Table 1 and Fig. S8), indicating that the TPase active side of PBP2^L61R^ is still accessible to mecillinam. Unfortunately, the co-expression of PBP2^L61R^ together with either MreC or MreD alone, or both together, was not possible in most cases, as FRET cells repeatedly lost the mKO-PBP2^L61R^ signal upon induction (Table 1), suggesting toxicity of these combinations. The cells did not lose the mKO signal in only two out of six attempts to co-express MreC and PBP2^L61R^. Of those samples the calculated Ef*_A_* value of the MreC and PBP2^L61R^ pair remained at 5.4 ± 1.7%, which was comparable with MreC-PBP2^WT^ (Fig. 4a and Table 1). These results suggest that the hyperactive mutant PBP2^L61R^ likely behaves similarly as wild type PBP2 in the interaction with its partner proteins.

**Figure 5.**
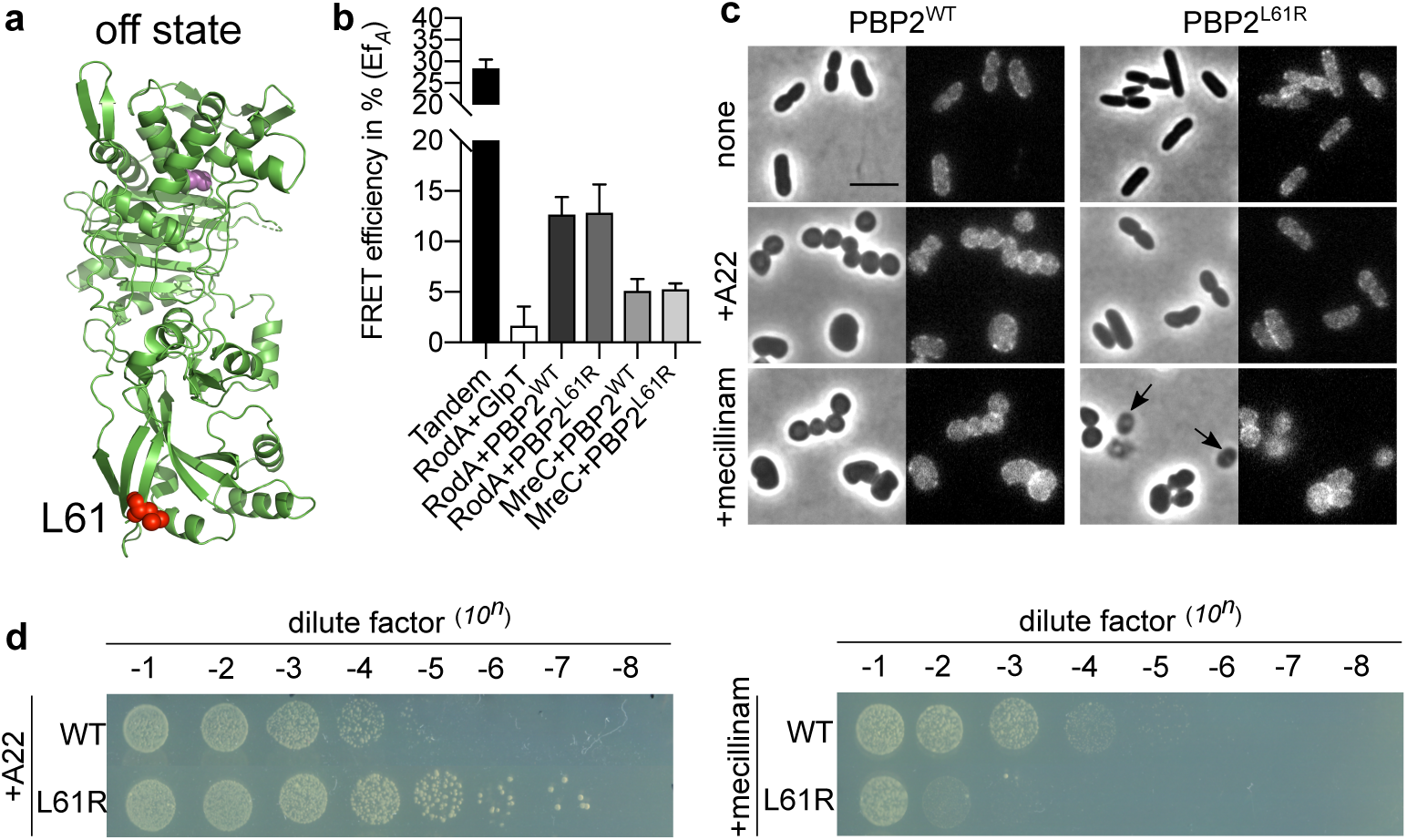
Hyperactive mutant PBP2^L61R^ is more sensitive to mecillinam. **a.** Modeled “off state” structure of *E. coli* PBP2 from *H. pylori* PBP2 using Phyre 2 (8). Structural information lacks on the juxta-membrane, transmembrane helix and cytoplasmic regions of PBP2. Residue Leu61 is colored in red and shown as spheres. Active site Ser330 is colored in pink and shown as spheres. **b**. Hyperactive mutant PBP2^L61R^ interacts similar like PBP2^WT^ with RodA and MreC. RodA and MreC were fused with mCherry, and PBP2^WT^ and PBP2^L61R^ were fused with mKO. **c**. Phase contrast and fluorescence images of cells expressing PBP2^L61R^ were less sensitive to A22 but hypersensitive to mecillinam in liquid culture. LMC500 strain expressing nothing, or PBP2^WT^, or PBP2^L61R^ were grown in LB medium at 37 °C. IPTG induction (15 μM), and A22 treatment (10 mg·L^-1^) or mecillinam treatment (2 mg·L^-1^) were applied to each culture for 2 mass doublings. Arrows indicate cells that lysed after mecillinam treatment in the PBP2^L61R^ culture. Scale bar equals 5 μm. **d**. Spot assay to test the sensitivities of PBP2^WT^ and PBP2^L61R^ to A22 (10 mg·L^-1^) and mecillinam (2 mg·L^-1^).

Having observed these unexpected results, we continued to further characterize the hyperactive PBP2^L61R^. Consistent with its reported functionality (51), we observed that mKO-PBP2^L61R^ was capable to support growth of the PBP2(TS) strain LMC582 at the non-permissive temperature (Supplementary Fig. 9b). However, the expression of mKO-PBP2^L61R^ resulted in longer and thinner cells (Fig. 5c and Supplementary Fig. 9c), and in reduced sensitivity of cells to the MreB inhibitor A22 (Fig. 5c) as reported (51). Interestingly, cells expressing PBP2^L61R^ were hypersensitive to mecillinam (Fig. 5c and d). These results indicate potential defects in the peptidoglycan layer of cells expressing PBP2^L61R^, and these defects may be tolerable under undisturbed growth conditions but are exacerbated in the presences of mecillinam. Considering that PBP2^L61R^ stimulates the GTase activity of RodA *in vitro* (51) and our results on the cellular interactions, it was possible that the L61R exchange in PBP2 enhances only the activity of RodA and has no effect on PBP2’s TPase activity. *In vitro* peptidoglycan synthesis experiments showed that inactivation of PBP2 by mecillinam resulted in longer glycan chains synthesized by a PBP1A-PBP2 complex (18), hence we wondered whether the presence of PBP2^L61R^ affected peptidoglycan synthesis in the cell. We prepared peptidoglycan and analyzed its composition from cells expressing PBP2^L61R^, wild type PBP2, and the control membrane protein GlpT. As predicted, the peptidoglycan from all strains retained a similar extent of peptide cross-linkage. By contrast, only the peptidoglycan from the PBP2^L61R^ expressing cells contained unusually long glycan chains with a mean length ∼52 disaccharide units (Table 3). The peptidoglycan of the strain overexpressing wild-type PBP2 had a mean glycan chain length of ∼38 units, and the mean glycan chain lengths of the other strains were between 40-43 disaccharide units (Table 3, Supplementary Table 3). A stimulating effect of PBP2^L61R^ on RodA’s GTase activity would be consistent with the previously observed A234T mutation in RodA that suppressed the morphological defects of MreC mutants (51), and would also explain why PBP2^L61R^ could only poorly restore survival and rod-shape in cells depleted of MreCD or RodZ (51). The tolerance to A22 and changes in MreB dynamics in the PBP2^L61R^ background (51) could also be explained by the enhanced RodA GTase activity by PBP2^L61R^; a direct interaction between RodA and MreB was detected with a Ef*_A_* value of 5.5% ± 1.7% (Table 1 and Supplementary Fig. 10). All together these results indicate that the ‘hyperactive’ PBP2^L61R^ likely has unchanged TPase activity itself but is probably in the off state conformation in the absence of MreC (44). The changes in cell morphology, the resistance to A22 and sensitivity to mecillinam, the partial compensation of MreCD-RodZ depletion and MreB dynamic changes are all likely due to an enhanced GTase activity of RodA (and perhaps PBP1A) (18, 51), which results into longer glycan chains in the peptidoglycan mesh.

**Table 3.** Summary of muropeptide composition of LMC500 strains carrying no plasmid or different expression plasmids.

| Muropeptides<sup>2</sup> or feature | Percent peak area (%)<sup>1</sup> |  |  |  |  |
| --- | --- | --- | --- | --- | --- |
|  | LMC500 | LMC500 mKO | LMC500 mKO-PBP2<sup>WT</sup> | LMC500 mKO-PBP2<sup>L61R</sup> | LMC500 mKO-GlpT |
| Monomers (total) | 52.6 ± 0.1 | 52.0 ± 0.0 | 52.1 ± 1.1 | 52.2 ± 0.9 | 52.2 ± 0.0 |
| dipeptides | 1.4 ± 0.0 | 1.6 ± 0.1 | 1.3 ± 0.1 | 1.2 ± 0.0 | 1.4 ± 0.0 |
| tripeptides | 5.0 ± 0.2 | 5.0 ± 0.0 | 5.0 ± 0.5 | 4.0 ± 0.3 | 4.7 ± 0.1 |
| tetrapeptides | 41.6 ± 0.0 | 41.0 ± 0.0 | 41.3 ± 0.5 | 43.3 ± 2.0 | 41.8 ± 0.2 |
| anhydro | 0.9 ± 0.0 | 1.0 ± 0.0 | 1.0 ± 0.0 | 0.7 ± 0.0 | 1.0 ± 0.0 |
| LysArg | 3.7 ± 0.4 | 3.4 ± 0.2 | 3.5 ± 0.5 | 3.0 ± 1.2 | 3.4 ± 0.5 |
| Dimers (total) | 40.5 ± 0.0 | 40.8 ± 0.0 | 40.2 ± 0.7 | 41.7 ± 0.9 | 40.7 ± 0.1 |
| tetratripeptide | 3.1 ± 0.3 | 3.0 ± 0.2 | 3.0 ± 0.1 | 2.2 ± 0.0 | 2.9 ± 0.1 |
| tetratetrapeptide | 36.0 ± 1.0 | 36.3 ± 0.2 | 35.0 ± 2.6 | 37.1 ± 2.7 | 36.0 ± 0.9 |
| anhydro | 2.0 ± 0.0 | 2.1 ± 0.0 | 2.2 ± 0.0 | 1.7 ± 0.0 | 2.0 ± 0.0 |
| LysArg | 1.6 ± 0.0 | 1.7 ± 0.1 | 1.5 ± 0.2 | 1.2 ± 0.2 | 1.6 ± 0.1 |
| Trimers (Total) | 4.4 ± 0.1 | 4.6 ± 0.1 | 4.8 ± 0.0 | 3.8 ± 0.0 | 4.5 ± 0.1 |
| Dipeptides (total) | 1.4 ± 0.0 | 1.6 ± 0.1 | 1.3 ± 0.1 | 1.2 ± 0.0 | 1.4 ± 0.0 |
| Tripeptides (total) | 7.2 ± 0.9 | 7.2 ± 0.2 | 7.6 ± 1.1 | 6.3 ± 0.7 | 7.0 ± 0.3 |
| Tetrapeptides (total) | 85.8 ± 0.1 | 85.8 ± 0.0 | 85.3 ± 0.1 | 87.7 ± 0.0 | 86.2 ± 0.0 |
| Chain ends (anhydro) | 2.3 ± 0.0 | 2.4 ± 0.0 | 2.5 ± 0.0 | 1.9 ± 0.0 | 2.3 ± 0.0 |
| Glycan chain length (DS)<sup>3</sup> | 42.8 ± 0.2 | 40.5 ± 0.0 | 38.3 ± 0.1 | **52.1 ± 0.5** | 41.6 ± 0.0 |
| Degree of cross-linkage | 23.2 ± 0.1 | 23.4 ± 0.0 | 23.3 ± 0.1 | 23.3 ± 0.2 | 23.3 ± 0.0 |
| % Peptides in cross-links | 47.4 ± 0.1 | 48.0 ± 0.0 | 48.0 ± 1.1 | 47.8 ± 0.9 | 47.8 ± 0.0 |

values are mean ± variation of two biological replicates.
<sup>2</sup> muropeptide names according to Glauner, 1988 (63).
<sup>3</sup> average glycan chain length in disaccharide (DS) units calculated from the percentage of anhydro-MurNAc containing muropeptides. The bold number highlights the increased average glycan chain length in cells expressing PBP2<sup>L61R</sup>.

## Discussion

### Peptidoglycan synthesis by the elongasome and divisome

In this work we aimed to reveal the regulation of peptidoglycan synthesis during length growth. Recent studies revealed a possible mechanism for the regulation of septal peptidoglycan synthesis by FtsBLQ and FtsN (19–21, 52–54). In this model, the FtsBLQ subcomplex inhibits the activities of PBP3 (consequently also inhibiting FtsW) and PBP1B, and keeps septal peptidoglycan synthesis in check. A small amount of FtsN is already present at pre-septal sites together with ZipA and the class A PBPs, PBP1A and PBP1B (55). However, only once FtsN accumulates at higher levels it is able to relieve the suppression of FtsBLQ on the peptidoglycan synthases, thereby activating septal peptidoglycan synthesis. During length growth, the elongasome proteins, such as MreC, MreD, PBP2, RodA and RodZ, localize in the lateral membrane, which makes it a challenge to investigate their cellular dynamics. Thus, it is still largely unknown how the elongasome regulates and coordinates peptidoglycan synthesis.

### MreCD proteins regulate the interaction between RodA and PBP2

In this study, we showed that RodA and PBP2 form a subcomplex independent of their biochemical activities (Fig. 2). This interaction requires the transmembrane helix and periplasmic parts of PBP2 (Fig. 3). *In vivo* FRET experiments revealed that MreC interacts with directly with PBP2, which modulated the interaction between PBP2 and RodA (Fig. 4). Surprisingly, MreD also interacts with PBP2 and but has an opposite effect, as it reverses the PBP2-RodA interaction change stimulated by MreC (Fig. 4). This is similar to the regulation of septal peptidoglycan synthesis where FtsN interacts with itself and accumulates at midcell (56, 57), and this accumulation is assumed to abolish the suppression of the peptidoglycan synthesis complex FtsW-PBP3-PBP1B by FtsBLQ. When comparing the cellular numbers of these proteins synthesized per generation (46), we noticed that the average number of FtsN molecules per cell is about 2 times higher than that of FtsBLQ and FtsW-PBP3 proteins. The number of MreC molecules is also about 2 times higher than that of MreD and PBP2-RodA molecules (46). MreC, but not MreD, is also reported to interact with itself, and the structural data showed that two molecules of MreC bind to one PBP2 molecule (44). Together with these published data, our results indicate that the balance between the MreC and MreD determines the nature of the interaction between PBP2 and RodA. Structural data show that the interaction between MreC and PBP2 causes a conformational change in PBP2 that was suggested to correspond to its activation from the off state to the on state (44). This conformational change could correspond to the change in the interaction between PBP2 and RodA induced by MreC. And likely, when MreD is co-expressed with MreC, the reversed change in the interaction between PBP2 and RodA could correspond to the PBP2 conformational change from the on state to the off state. This potential of MreC and MreD to affect the RodA-PBP2 interaction could reflect the regulation of elongasome activity and peptidoglycan synthesis during length growth. The overexpression of either MreC or MreD would shift this balance and result in over activation or suppression of PBP2-RodA activities, and cause morphological defects (Fig. 4e and Supplementary Fig. 7).

Based on our observations, we propose a model for the regulation of PBP2 (elongasome) activity and cylindrical peptidoglycan synthesis (Fig. 6). The peptidoglycan synthases RodA and PBP2 form a stable subcomplex. MreC stimulates and activates PBP2 and RodA, while MreD interferes with the PBP2 MreC interaction to keep PBP2 activity and peptidoglycan synthesis in check. The further binding and accumulation of MreC to PPB2 would eventually outcompete MreD, which will activate PBP2 and consequently initiate peptidoglycan synthesis. Because hydrolytic activity is required to allow insertion of newly synthesized peptidoglycan into the existing mesh (58, 59), a balanced regulation is likely needed to avoid premature glycan strand synthesis. Consequently, the elonsasome, like the divisome, must have a mechanism to sense whether all partners are at the right position to act. A well-regulated moment for the switching on of peptidoglycan synthesis by balancing the MreCD ration is likely part of such a regulatory system.

**Figure 6.**
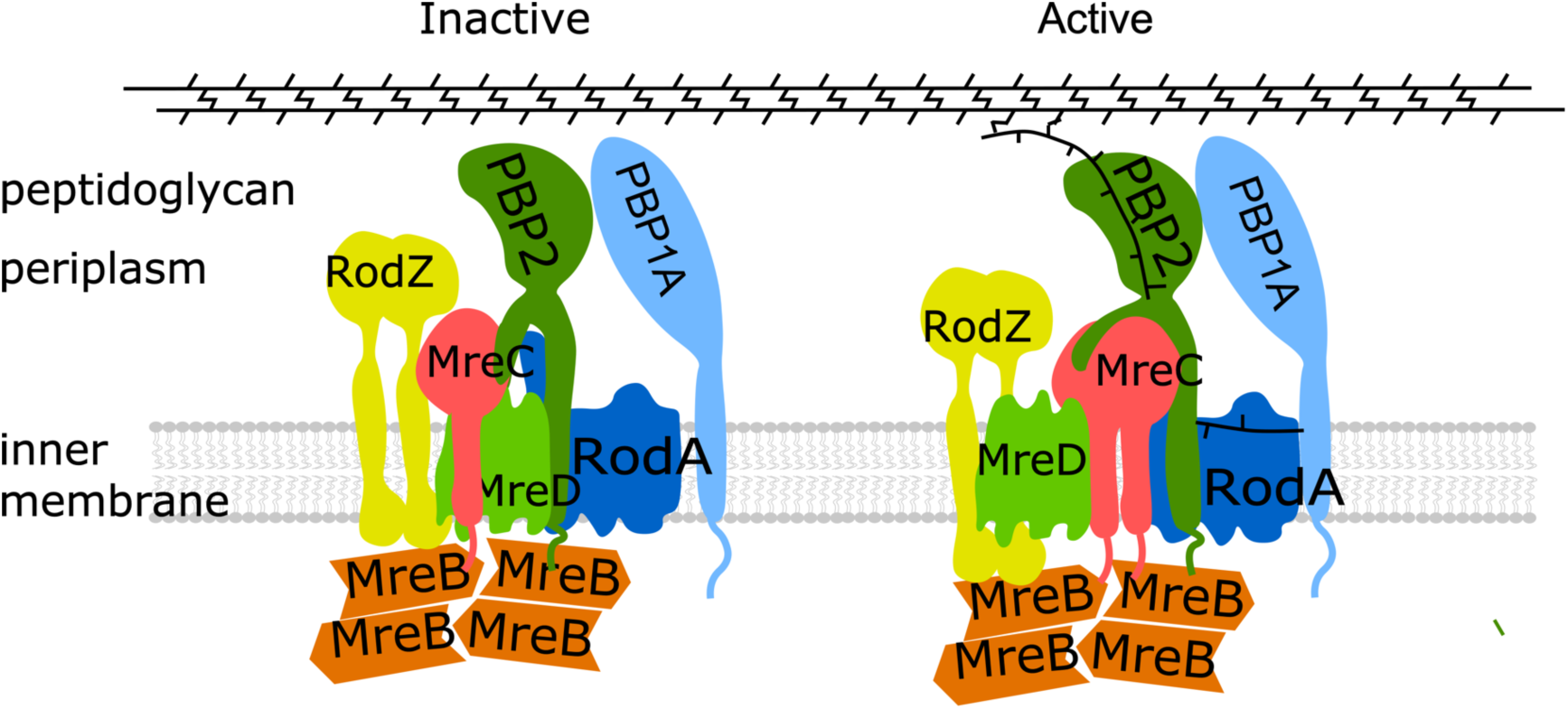
Model for regulation of elongasome and PG synthesis. RodA and PBP2 interact with each other and form a stable subcomplex. MreC and RodZ interact strongly with MreB filaments that likely link MreB to the PG synthesis proteins. MreC interacts with PBP2 that could stimulate and activate PBP2, while MreD, which interacts with both PBP2 and MreC, suppresses the activation of PBP2 by MreC, and keeps PG synthesis under control. The accumulation of MreC to the elongasome will finally abolish the inhibition of MreD and activate PBP2 by changing its conformation from the “off state” to the “on state”, and subsequently activate the elongasome and PG synthesis.

## Materials and Methods

### Media, strains, plasmids and primers

LB (10 g tryptone, 5 g yeast extract and 10 g NaCl, per liter) and Gb4 (6.33g K_2_HPO_4_·3H_2_O, 2.95g KH_2_PO_4_, 1.05 g (NH_4_)_2_SO_4_, 0.10 g MgSO_4_·7H_2_O, 0.28 mg FeSO_4_·7H_2_O, 7.1 mg Ca(NO_3_)_2_·4H_2_O, 4 mg thiamine, 2 mg uracil, 2 mg lysine, 2 mg thymine, and 0.5 % glucose, per liter, pH 7.0) were used for cell cultures in rich and minimal medium, respectively, as indicated. Final concentrations for antibiotics were: 100 μg·L^-1^ ampicillin, 50 μg·L^-1^ kanamycin and 25 μg·L^-1^ chloramphenicol.

*E. coli* strains and plasmids used in this study are listed in supplementary Table 1. Primers used in this study were listed in supplementary Table 2. The plasmids were constructed as following:

***pXL29***. Plasmid pXL28 and pWA004 were digested with *EcoR*I and *Hind*III restriction enzymes, the generated (GGS)_2_-GlpT expressing gene and pSAV057-mKO linear vector were ligated together to generate the mKO-(GGS)_2_-GlpT expressing plasmid.

***pXL36, pXL40, pXL44, pXL48, pXL56 and pXL63***. Plasmids pXL36 and pXL40 that expressing mCherry-fused RodA^R109A^ and RodA^Q207R^ were generated from pSAV047-RodA by mutagenesis PCR using primer pairs priXL61-priXL61 and priXL69-priXL70, respectively. To construct non-fused version of RodA variants, wild type *rodA* gene was amplified using primer priXL59 and priXL60 from the MG1655 genomic DNA and ligated into empty pSAV057 vector, to generate plasmid pXL63. The two mutants plasmids were generated in the same way as descripted above from pXL63. mKO fused RodA plasmid pXL56 was constructed by cutting and pasting the *rodA* gene from pSAV047-RodA to the pSAV058 plasmid with *EcoRI* and *HindIII* restriction enzymes.

***pXL148, pXL149, pXL158 and pXL159***. The PBP2 domain swap plasmids were constructed by Gibson assembly (60). For N-terminus replacement, primer pairs priXL146-priXL258 and priXL147-priXL259 were used for PCR from plasmid pWA004. The PCR products were purified after *DpnI* digestion and assembled to generated pXL148 that excludes the N-terminus of PBP2. For pXL149, the primer pair priXL258-priXL260 was used to amplify the entire pWA004 plasmid excluding the first 45 residues. Primer pair priXL261 and priXL263 was use to amplify the first 37 residues of MalF from MG1655 genome. The PCR products were purified after *DpnI* digestion and assembled to generate pXL149 plasmid. The PBP2^S330C^ and PBP2^L61R^ plasmids were constructed with mutagenesis PCR from the pWA004 plasmid using primer pairs priXL274-priXL275 and priXL276-priXL277, respectively.

***pXL165***, ***pXL166 and pXL169***. *mreC*, *mreCD* and *mreD* genes were amplified from MG1655 genome using primer pairs priXL282-priXL286, priXL282-priXL283 and priXL299-priXL283, respectively, and cloned into plasmid pSAV047 with *EcoRI* and *HindIII* restriction enzymes, to generate the mCherry fused version of these genes.

***pXL167 and pXL168***. The third plasmids used in the three-plasmids FRET experiments. *mreC* and *mreCD* genes were cloned into plasmid pSG4K5 (61) with Gibson assembly, respectively, and the p*_trcdown_* promoter was introduced to control the protein expression. Primer pair priXL294-priXL295 was used to amplify the linear vector from pSG4K5. Primer pair priXL296-pp15 was used to amplify the p*_trcdown_* promoter. Primer pairs pXL284-priXL297 and priXL284-priXL298 were used to amplify the *mreCD* and *mreD* genes, respectively.

### Bacterial growth, morphology and protein localization

For general growth experiments in rich medium, overnight cultures (37 °C) were diluted 1:1000 into fresh LB medium with 0.5% glucose and the required antibiotics, and grew to OD_600_ around 0.2 at 37 °C. Cultures were further diluted 1:5 into fresh LB medium with required antibiotics, and induced with 15 μM IPTG for 2 mass doubling at 37 °C (OD_600_ reached around 0.2).

For complementation experiments, temperature sensitive strains expressing the mutant plasmids were grown as described above at 30 °C. Cultures were further diluted 1:5 into fresh LB medium with required antibiotics, and induced with 15 μM IPTG for 2 mass doubling at 30 °C and 42 °C, respectively (OD_600_ reached around 0.2).

After induction, cells were fixed with FAGA (2.8 % formaldehyde and 0.04 % glutaraldehyde, final concentration) for 15 minutes and centrifuged at 7000 rpm for 10 min at room temperature. Cell pellets were suspended and washed 3 times with PBS (pH7.2) buffer. Subsequently, bacterial morphology and protein localization were imaged by wide field phase contrast and fluorescence microscopy. Specially, cells expressing the mKO fused proteins were firstly matured at 37 °C overnight before imaging by microscopy.

### FRET experiment and data analysis

Protein interactions were detected by FRET as described previously (33, 37, 62). For the FRET experiments, mCherry and mKO fluorescent proteins were used as acceptor and donor fluorophores, respectively. LMC500 strain was co-transformed with the FRET pairs that were to be detected. In each FRET experiment, the empty-vector reference, mCherry reference, mKO reference were included to be able to calculate the Ef*_A_* by unmixing of the measured FRET pair spectrum in its individual components; background, mCherry, mKO and sensitized emission spectra. A tandem fusion of mKO-mCherry was used as positive control, and the mCherry-RodA and mKO-GlpT pair was used as negative control. After transformation, FRET strains were firstly grown in LB medium (with antibiotics and 0.5% glucose) overnight at 37 °C, and diluted 1:1000 into fresh medium and grown to OD_600_ around 0.2 at 37 °C. Subsequently, FRET strains were diluted 1:500 into Gb4 medium and grown to steady state at 28 °C (OD_450_ was kept below 0.2). All FRET strains were induced with 15 μM IPTG (and treated with mecillinam at 2 mg·L^-1^ concentration as indicated) for two mass doubling before FAGA fixation. After fixation, FRET cells were pelleted by centrifugation at 7000rpm at room temperature and washed 3 times with PBS buffer (pH 7.2). Then all samples were incubated at 37 °C overnight and stored at 4 °C for 1 extra day before measured with spectrofluorimeter (Photon Technology International, NJ). Emission spectra of acceptor and donor fluorophores were measured through 6-nm slit widths with 1 second integration time per scanned nm for 3 times averaging. Filters 587/11 nm (587/11 nm BrightLine single band-pass filter, Semrock, New York, NY, USA) and 600nm long-pass (LP) filter (Chroma Technology Corp., Bellows Falls, VT) were used for excitation and emission of acceptor fluorophore (mCherry), while 541/12 nm (Semrock) and 550 nm long pass (Chroma) filters were used for mKO excitation and emission, respectively. For calculation, measurement of PBS buffer was subtracted from all samples, and the empty-cell reference was subtracted from the donor and acceptor spectra. The FRET efficiencies were calculated as described previously (37, 62).

For three plasmids FRET, a third plasmid (expressing MreC or expressing MreCD both) was introduced into the whole two plasmids FRET system. Empty pSG4K5 vector was also introduced as a control to correct for the reduction in FRET efficiency due to the burden of maintaining three plasmids.

### Spot assay

To test the sensitivity of *E. coli* strains to A22 and mecillinam, LMC500 strain was transformed with pWA004 (PBP2^WT^) or pXL159 (PBP2^L61R^). Strains expressing each construct were grown in LB medium as descried above without induction. Cell cultures were diluted with varying dilution factors (Fig. 4C). A drop of 10 μl cell culture from each dilution was loaded on the LB agar dish (with chloramphenicol, 15 μM IPTG, 10 μg·mL^-1^ A22 or 2 μg·mL^-1^ mecillinam) and incubated overnight at 37 °C.

### Peptidoglycan analysis

Peptidoglycan sacculi were prepared from *E. coli* cells, digested with cellosyl (kind gift from Hoechst, Germany), reduced with sodium borohydride and analyzed by high pressure liquid chromatography as described (Separation and quantification of muropeptides with high-performance liquid chromatography) (63).

### Microscopy

Bacterial cell samples were immobilized on 1.3% agarose pads (w/v in Gb4 medium) and imaged under microscopy. Fluorescence microscopy was carried out either with an Olympus BX-60 fluorescence microscope equipped with an UPlanApo 100×/N.A. 1.35 oil Iris Ph3 objective, or with a Nikon Eclipse Ti microscope equipped with a C11440-22CU Hamamatsu ORCA camera, a CFI Plan Apochromat DM 100× oil objective, an Intensilight HG 130W lamp and the NIS elements software (version 4.20.01). Images were acquired using the Micro Manager 1.4 plugin for ImageJ, and analyzed with Coli-Inspector supported by the ObjectJ plugin for ImageJ (version 1.49v) (64).

## Acknowledgements

X.L. was supported by the Chinese Scholarship Council (File No.201406220123). W.V. received funding from the Wellcome Trust (101824/Z/13/Z). We thank Lisa Atkinson for preparation of peptidoglycan sacculi and Prof. Thomas Bernhardt for the gift of modeled *E. coli* PBP2 structures(51).

## Author contributions

X.L and T.B. designed and analyzed all the experiments. X.L. performed all experiments. Jacob Biboy and Waldemar Vollmer performed the peptidoglycan analysis experiment and analyzed these data. X.L. and T.B. analyzed all FRET data and wrote the manuscript.

